# Pain reflects the informational value of nociceptive inputs

**DOI:** 10.1101/2023.07.14.549006

**Authors:** Michel-Pierre Coll, Zoey Walden, Pierre-Alexandre Bourgoin, Véronique Taylor, Pierre Rainville, Manon Robert, Dang Khoa Nguyen, Pierre Jolicoeur, Mathieu Roy

**Author notes:** Correspondance to: Mathieu Roy. Department of Psychology, McGill University, 2001 McGill College, Montréal, QC, Canada. https://www.mcgill.ca/psychology/mathieu-roy.

## Abstract

Pain perception and its modulation are fundamental to human learning and adaptive behavior. This study investigated the hypothesis that pain perception is tied to pain’s learning function. Thirty-one participants performed a threat conditioning task where certain cues were associated with a possibility of receiving a painful electric shock. The cues that signalled potential pain or safety were regularly changed, requiring participants to continually establish new associations. Using computational models, we quantified participants’ pain expectations and prediction errors throughout the task and assessed their relationship with pain perception and electrophysiological responses. Our findings suggest that subjective pain perception increases with prediction error, that is when pain was unexpected. Prediction errors were also related to physiological nociceptive responses, including the amplitude of the nociceptive flexion reflex and EEG markers of cortical nociceptive processing (N2-P2 evoked potential and gamma-band power). Additionally, higher pain expectations were related to increased late event-related potential responses and alpha/beta decreases in amplitude during cue presentation. These results further strengthen the idea of a crucial link between pain and learning and suggest that understanding the influence of learning mechanisms in pain modulation could help us understand when and why pain perception is modulated in health and disease.

## Introduction

Pain not only serves as a warning signal, but it also teaches us to identify and avoid future threats [50]. This intimate link between pain and learning is supported by studies showing that the prediction error (PE) signals necessary to learn from pain are linked to activity in cerebral targets of ascending nociceptive pathways [24,29,48], as well as to changes in gamma oscillations in response to surprising pain [8,55]. However, it remains unclear if the learning processes related to pain occur separately from the processes associated with subjectively perceived pain [8] or if perceived pain is more deeply tied to its learning function [17,55].

In a previous study, we demonstrated that the amplitude of the spinally-mediated nociceptive flexion reflex (NFR) and perceived pain were associated with learning parameters during a threat conditioning procedure [57]. These findings suggested that the nociceptive signals that originate from the spinal cord and that ultimately produce our subjective pain perception may have a learning function. However, the low number of changes in cue-outcome contingencies in this study made it difficult to examine PEs specifically. Another problem with that interpretation is that the amplitude of the NFR does not necessarily index the excitability of spinal nociceptive neurons because the reflex’s motor output can be modulated independently from its sensory input [23]. To overcome these limitations, we designed a study in which the reinforcement contingencies changed rapidly so as to produce frequent PEs and combined the recording of NFR amplitude with electroencephalography (EEG) to link NFR modulation with ascending nociceptive signals to the cortex. We hypothesized that a positive correlation between NFR amplitudes and pain-related EEG signals and pain ratings would demonstrate that pain’s learning and sensory aspects are tied together at the spinal level.

Early somatosensory-evoked potentials, such as the N2-P2, are produced by ascending nociceptive signals to the cortex. More specifically, the N2-P2 is thought to originate from midline generators linked to pain’s affective-motivational dimension [18] and has been correlated with subjective pain reports [11,19], although it may also reflect non-pain-specific aspects of somatosensory processing [27,40]. By contrast, gamma band activity has recently been suggested as a more specific indicator of pain [26,35] and can be used to complement the N2-P2 as a cortical index of pain processing.

Finally, to understand how PEs are generated, it is important to consider the cerebral processes associated with the expected probability of pain before it occurs. The late positive evoked potential (LPP) and a decrease of oscillatory power in the alpha and beta bands [41,43] have both been linked with expectation of aversive events and pain perception [2,41,55], but their link to computational estimates of learning such as PEs have yet to be established. Given the intimate link between perception and PE across modalities [15,25,54], we hypothesized that that PE estimates would correlate with NFR amplitude, N2-P2 amplitude, gamma oscillations and pain ratings in response to the nociceptive stimulus and that computational estimates of increased pain probability would be associated with an increased LPP and decreased alpha-beta power during pain anticipation.

## Materials and Methods

### Participants

Thirty-five healthy participants with no history of neurological or pain disorders took part in the experiment. Four participants were excluded from all analyses. One participant exhibited no discernible skin conductance responses (SCR). Another participant’s skin conductance data contained extensive flat segments due to technical problems. Additionally, two participants had fewer than 20 clean trials remaining in each condition of the EEG data following the cleaning procedure detailed below. All analyses were thus performed on 31 participants (11 males, mean age: 23.39 years, SD=1.83). All participants provided written informed consent, and the experimental procedures were approved by the local ethics committee (*Comité mixte d’éthique de la recherche du Regroupement Neuroimagerie Québec* - Project #CMER RNQ 11-12-014).

### Stimuli

#### Visual Stimuli

Fourteen still frames showing neutral faces from different individuals (7 females) were selected from the Radboud Faces Database [32], were grey-scaled and cropped to remove non-facial features and served as the conditioned stimuli (CS). The stimuli-condition associations and presentation order were fully randomized.

#### Electrical Stimulations

The unconditioned stimulus (US) was presented simultaneously with the offset of the CS and consisted of a 30 ms transcutaneous electrical shock (10 x 1ms pulses at 333 Hz) delivered with a Grass s48 pulse stimulator triggered by a computer running the E-Prime 2 software (Psychology Software Tools, Inc., Pittsburgh, USA). The electric shock was delivered to alcohol-cleaned skin over the right sural nerve through a custom-made pair of 1-cm^2^ bipolar stimulating electrodes placed behind the malleus of the right ankle with a distance of 2 cm between the anode and the cathode.

The shock intensity was calibrated to each participant’s sensitivity using a staircase procedure [61] used in previous studies (e.g. [56,57]). In this procedure, stimulations were delivered from the lowest level of intensity and incremented in steps of 0.5 or 1mA at intervals of 6s. After each stimulation, participants reported if they perceived the stimulation and provided a pain rating on a 0 to 100 scale, with 0 being no pain and 100 being the worst pain imaginable. The lowest level of stimulation detectable by the participant was determined as the detection threshold, and the lowest level of intensity perceived as painful was determined as the pain threshold. Once levels of tolerance were reached (the highest level of intensity considered acceptable for the participant) or distinguishable muscular electromyographic (EMG) activity was elicited from baseline EMG in the NFR response window (90-180ms) for at least 3 consecutive stimulations, stimulations were administered in decrements of 1mA until the lowest level of administration. This procedure was repeated three times. A stimulus–response graph was then plotted to display NFR amplitudes (the integral of the root-mean-square transform of the EMG between 90-180 ms) by stimulus intensity. The NFR threshold was determined as the intensity for which 50% of NFR scores were higher than baseline noise-related EMG activity levels, and a stimulus intensity corresponding to 135% of the NFR threshold was selected. Finally, ten stimulations were administered at this intensity in order to ensure that the NFR was consistently elicited and to account for any habituation effects. Participants were then asked to provide a pain intensity rating for these stimulations. If the stimulation led to perceived pain, was considered to be tolerable and led to at least eight visible NFRs, it was selected as the US for the conditioning experiment. Otherwise, the intensity was slightly increased or decreased until a painful but tolerable stimulation eliciting reliable NFR was obtained and then used for the conditioning task. The mean selected stimulation intensity was 8.35 mA (SD=3.66) and the mean pain rating for this stimulation was 38.03 (SD= 26.05) out of 100.

### Procedure

Throughout the experiment, participants sat in an electrically shielded room. After the recording electrodes were placed, the pain thresholding procedure was performed to determine the intensity of the electric shock that should be used for the task. Once the stimulation intensity was selected, the EEG cap was set up.

Participants were told they would see faces that may be followed by electric shocks and that the relationship between faces and shocks could change over time. They were instructed to look at the fixation cross and faces at all times. Finally, participants were asked to evaluate the pain induced by electrical stimulations using a visual analogue scale with the labels “No pain” and “Worst pain imaginable” at each extremity (0 no pain to 100 worst pain imaginable). The scale was presented as a horizontal bar on a computer screen, with the subject moving a cursor on the scale using a keyboard.

Participants then underwent the threat conditioning paradigm (see Figure 1a-b). The first block, represented by the gray column on the left side of Figure 1a, consisted of a habituation phase in which two faces are presented without any shocks. In the other six blocks, one of the 4 presented faces is co-terminated with a shock for 50% of the trials and the other faces are never paired with the shock (CS-). Each presentation of the CS+ face was followed by a fixation cross of 2s, and then the subjects were given a maximum of 10s to record their pain rating and to allow for the recording of the SCR. All trials started with an interstimulus interval (ISI) that was jittered between 5 to 7 seconds.

**Figure 1.**
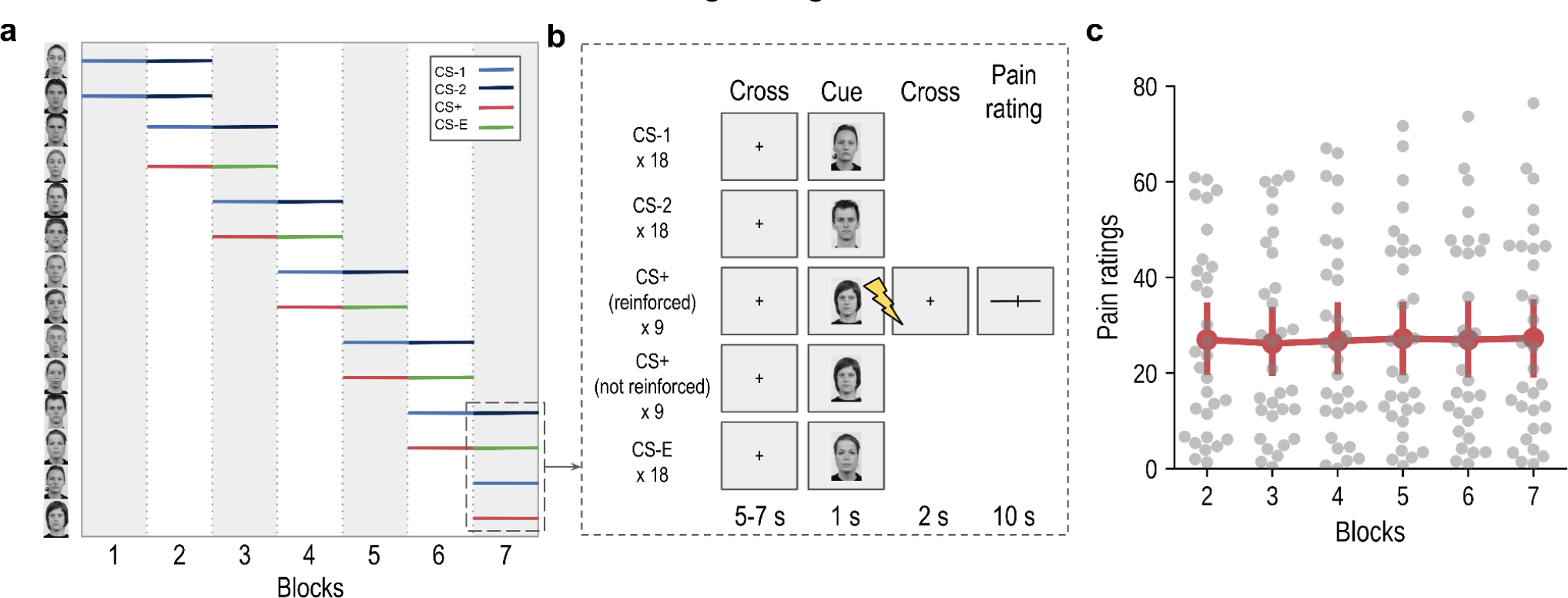
Illustration of the structure of the threat conditioning task and obtained pain intensity ratings. **(a)** The experiment began with the random presentation of two faces unpaired with electric shocks. After 18 presentations of each face, two new faces were introduced, one paired with shocks on 50% of trials (CS+), and one unpaired (CS-1), while the two initial faces were presented an additional 18 times (CS-2). Then, as the two initial faces stopped being presented, two new faces were introduced, one paired with shocks, and one unpaired, and the previously paired face became unpaired (CS-E). Each face was presented a total of 36 times, except for the last two faces, each presented 18 times. **(b)** I**llustration of the trial structure within blocks 2 to 7**. For each trial, a fixation cross was first presented for 5 to 7 s, followed by the CS for 1 s. Shocks were delivered at the offset of 50% of the CS+, followed by a fixation cross (2s) and a visual analog scale (VAS) displayed for pain ratings (10s). Only trials in which a shock was delivered (CS+ reinforced) were followed by a rating scale for a total of 54 ratings during the experiment. **(c)** Average pain intensity ratings (red) provided across the experimental blocs with reinforced cues and dots showing the average rating of each participant for each block. Error bars show the 95% confidence interval.

The presentation of the stimuli was divided into 7 blocks. Each face was presented 18 times in a single block for a total of 36 presentations of each face except the two last ones that were only presented for one block (18 times). Two new faces were presented in each block, with one of the new faces becoming the new CS+ face while the other remained CS-. CS+ faces in one block systematically became a CS-extinct (CS-E) face in the next block. The trials were presented in pseudo-random order, with the constraint that the first CS+ face of each block was always reinforced with a shock.

## Physiological measures

### Electrodermal recordings and processing

SCR were recorded with a BIOPAC MP160 system combined with an EDA100C module using a pair of gelled electrodes positioned on the palm of the participant’s right hand. The signal was amplified (5 µs/volt), bandpass filtered (1-5 Hz) and sampled at 1000 Hz. SCRs were analyzed using the PsPM toolbox [3] within MATLAB (R2019a, The Mathworks Inc., Natick, MA). Skin conductance data were first preprocessed using PsPM’s quality assurance script. PsPM’s default values were used to remove and interpolate data points above or below 60 or 0.05 µS, respectively or data points with a difference exceeding a slope of 10 µS/s. The data were then resampled to 10 Hz and normalized before analysis. SCR to each stimulus was estimated using a general linear model (GLM) approach by convolving a standard canonical SCR basis function with a time derivative to event onsets. We built a GLM for each trial which included the onset of the cue of the modeled trial as the regressor of interest and the onsets of the other cues, fixation crosses, shocks, rating screens and the onset of the recording as events of no interest. SCR amplitudes to the face cues were extracted using the reconstructed SCR [4] and used to fit the computational models.

### EMG recordings and processing

EMG activity of the biceps femoris was recorded through a pair of electrodes placed on the short head of the right biceps femoris and a reference electrode placed on the tibial tuberosity. The skin regions were shaved, exfoliated and cleaned with alcohol before placing the electrodes. The EMG signal was amplified by a 1000 factor and sampled at 6250 Hz with a BIOPAC MP160 equipped with the EMG100C. The EMG signal was resampled to 1000 Hz and rectified using the root mean square (RMS) transform with a 15 ms time constant. Finally, the NFR amplitude was calculated as the RMS EMG area under the curve between 90 to 180 ms after the end of the electric shock.

### EEG recordings and processing

EEG was recorded using 66 Ag-AgCl BioSemi Active Two electrode system (Biosemi Inc., Amsterdam, Netherlands) placed on the scalp according to the 10-10 international system. In Biosemi systems, a common mode sense and driven right leg electrodes placed on the cap serve as the ground, and all scalp electrodes were referenced to the common mode sense during recording. An electrooculogram (EOG) was recorded from 3 surface electrodes, two placed at the outer orbit of each eye horizontal to the pupil, and one placed below the left eye vertical to the pupil. Signals were amplified and digitized at a sampling rate of 1024 Hz. For 3 participants, the EEG signal was mistakenly recorded at 2048 Hz and was subsequently downsampled to 1024 Hz before preprocessing.

All EEG analyses were performed using MNE-python [21]. The data were first resampled to 1024 Hz if acquired at a higher rate and high-pass filtered at 0.1 Hz using a zero-phase non-causal finite impulse response (FIR) filter. Bad channels were identified via visual inspection and removed from the data mean number of bad channels: 1.61, SD=1.28, range=0-5). referenced to the average of all channels. An independent component analysis (ICA) [30] was performed using the *extended infomax* algorithm. Before ICA, the data were additionally bandpass filtered between 1 and 100 Hz, segmented −1 to 2 s around cues onsets (which included the shock when present), and segments with a value exceeding 500 uV were dropped before ICA. Artifactual components associated with eye blinks, eye movements, muscle artifacts, channel noise, line noise and heartbeats were first identified using the mne-ICAlabel algorithm [33] and validated manually. This led to the removal of an average of 11.35 components per participant (SD = 3.54, range=5-19). The remaining non-artifactual components were back-projected in the continuous data, which resulted in ICA-corrected data. After ICA, removed channels were interpolated using neighbouring channels, and only the ICA-corrected continuous data were used for subsequent analyses.

For event-related potentail (ERP) analyses, the ICA-corrected continuous data were additionally low-pass filtered at 30 Hz and the data was segmented −0.2 to 1 s around the onset of the face cue and −0.2 to 0.5 around the shock stimulation and baseline-corrected by subtracting the mean pre-stimulus value. Segments with values exceeding +/- 150 µV were excluded from further analyses, which led to the removal of an average of 9.09 % of trials (SD=7.18, range:0.64-28.41%) with a final average number of trials of 132.42 (SD=10.44, range:98-143) trials for the CS-1 condition, 116.39 (SD=8.94, range: 93-126) trials for CS-2, 93.25 (SD=12.42, range: 66-108) trials for CS+, 83.38 (SD=6.77, range: 63-90) trials for CS-E and 44.00 (SD=9.92, range: 20-54) trials for shocks. For the statistical analysis of ERPs in response to the shocks, we considered the amplitude of the N2-P2 evoked potential. The N2-P2 was measured at each trial for each participant as the difference between the peak amplitude between 200 and 500 ms and the preceding negative peak between 50 to 200 ms after the onset of the shock at electrode Cz.

For time-frequency representation (TFR) analyses, the ICA-corrected continuous EEG data segmented −2 to 2 s around the onset of the face cue or the shocks for the time-frequency analysis. To avoid capturing the response to the electrical shocks into the time-frequency response to the cue, only non-reinforced trials were analyzed for analyses of the time-frequency response to the visual cue unless otherwise specified. The same trials rejected from the ERP analyses using the approach described above were also removed from the time-frequency analyses for consistency. The TFR of each clean trial was obtained using a Morlet wavelet transform with a frequency range of 4 to 100 Hz and a resolution of 1 Hz and cycle sizes of 0.5*ƒ. The resulting TFR data were downsampled in time to 256 Hz, baseline corrected by taking the log of the ratio of each value and the −0.5 to −0.2 s pre-stimulus period and cropped to −0.2 to 1 s for subsequent statistical analyses. Based on visual inspection and previous studies [see 29], gamma-band power in response to the shock was measured as the mean power between 150 to 350 ms and 70 to 95 Hz after the onset of the shock at electrodes Cz, C1 and C2.

### Computational Modeling

We considered three learning models to explain the trial-wise anticipatory SCR responses: the Rescorla-Wagner model (RW), the Rescorla-Wagner/Pierce-Hall hybrid model (RW/PH) [44,46] and the Hierarchical Gaussian Filter (HGF; [38,39]). Two versions of each model were run. In the first version, the model was fitted for each cue without considering the influence of other cues (cue-specific). In a second version, the information about other seen cues is updated simultaneously when a cue is reinforced (inter-cue). Bayesian model comparison indicated that the inter-cue family of models showed an overall better fit to our data (exceedance probability of the intercue family: 0.90, model frequency: 61%) and only these models were further considered in addition to a null model in which no learning occurred. All models were fitted to participants’ individual trial-wise SCR using the TAPAS toolbox (https://github.com/translationalneuromodeling/tapas). Where priors were required for the HGF models, we selected them by inverting the perceptual model in isolation for each participant under suitably uninformative priors and used the group average of each parameter as prior. Trials that were reinforced or that were more than three deviations away from the participant’s mean SCR were ignored in the model fitting. The Log-model evidence (LME) was used to assess model fit using a random-effect Bayesian model selection implemented in the Variational Bayesian Analysis toolbox (VBA; https://mbb-team.github.io/VBA-toolbox; [14]).

### Null model

#### Perceptual model

We included a null model to ensure that our learning models would fit the data better than a model in which no learning occurred. The null model considered that participants could perfectly predict the occurence of the shock:

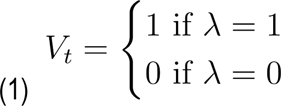

Where *V* is the expected probability and *λ* is the reinforcement value.

**Response model.** The response model for the null model considered that the SCR was linearly related to the perfect prediction of the shock:

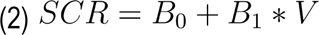

### Rescorla-Wagner model

#### Perceptual Model

The RW depends on prediction errors to direct associative changes, such that larger errors between the actual and expected value will drive larger changes in associative strength. The basic RW model is defined by equations 1 below where *V* is the expected shock probability which is updated every trial (t) according to the learning rate parameter *α*, the difference between the actual outcome and expected outcome (δ), *λ* the reinforcement value (shock = 1, no shock = 0), and *V* the expected probability.

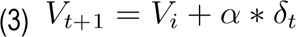

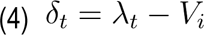

At the first presentation of each cue the expected probability corresponds to the fitted initial value, *V_i_* = *V_0_*. After each update, the updated expected value of the cue i is stored for future retrieval:

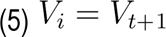

And cue i is added to the set of seen cues:

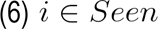

When a shock is delivered, each cue in the ensemble of all previously seen cues but not presented at trial t is updated using equations 1 and 2 with *λ* and the corresponding updated expected value is stored for each cue for future retrieval.

#### Response model

The RW response model considered that the SCR amplitude at each trial was linearly related to the expected value with the addition of Gaussian noise (*ζ*).

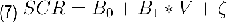

#### Rescorla-Wagner/Pearce-Hall Hybrid

##### Perceptual model

In the RW/PH model, the learning rate is influenced by the amount of attention devoted to the cue after a surprising outcome represented by an associability term. If there is a large error between the actual and expected outcome (e.g. a surprising event), this will result in an increased amount of attention being given to the relevant cues in order to learn associations; in contrast, small or no errors will hamper learning [44]. The hybrid model incorporates the PH into the RW model by updating reinforcement expectation by allowing for modulation of the learning rate (α) based on the degree of surprise regarding the event.

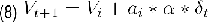

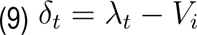

Where *t* is the trial number, *i* is the identity of the presented cue at trial t, *α* the learning rate, *δ* the prediction error, *λ* the reinforcement value, *V* the expected probability, *a* the associability and *γ* is a discount factor that determines how much *a* is influenced by events on the immediately preceding trials. At the first presentation of each cue, the expected probability and the associability correspond to the fitted initial value; *V_i_* = *V_0_*, *a_i_* = *a_0_*. After each update, the updated expected value and associability of cue i are stored for future retrieval:

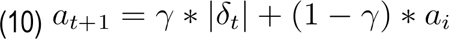

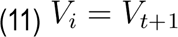

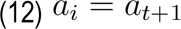

and cue i is added to the ensemble of seen cues (equation 3). When a shock is delivered, the expected value and associability of each previously seen cue but not presented at trial *t* is updated using equations 7, 8 and 9 with *λ* and the corresponding updated expected value and associability are stored for each cue.

##### Response model

Based on previous reports indicating that the SCR response is best characterized as a mixture of expectation and uncertainty, the RW/PH response model considered that SCR amplitude at each trial was linearly related to the expected value and associability with the addition of Gaussian noise.

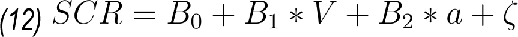

#### Hierarchical Gaussian Filter

##### Perceptual Model

The HGF consists of a perceptual and response model of an ideal Bayesian observer who is receiving inputs from the environment and has behavioural outputs (or responses, [39]). We used the standard two-level HGF for binary responses. As in the RW and RW/PH models described above, previously seen cues were updated with a negative prediction error when another cue was reinforced. Since the HGF was not found to outperform the RW model in the current experiment, for the sake of brevity, we refer the reader to previous descriptions of this model for the equations [38,39] and to the open code for details on the current implementation.

##### Response models

Similarly to the RW/PH model, the HGF response model included both the participants’ expectations and uncertainty at the first level:

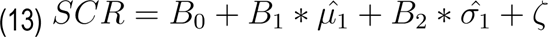

### Statistical analyses

#### Pain intensity ratings

Average pain intensity ratings collected on the visual analog scale were scored between 0 (no pain) and 100 (worst pain imaginable) and compared across blocks using a repeated-measures ANOVA to assess if there was a change in pain perception across blocks with reinforced cues (blocs 2 to 7). The ANOVA result was corrected using the Greenhouse-Geisser correction to take into account a violation of the sphericity assumption.

#### SCR

To confirm that conditioned cues elicited anticipatory SCR to threat-conditioned cues and to assess the habituation of SCR responses across the experiment, we used a linear mixed-model with the factor Cue type (CS+, CS-E, CS-1, CS-2), Block (2-7) and their interaction as fixed effects and participants as random effects allowing for random intercepts. The contribution of each fixed effect was assessed by comparing the models with and without this factor using a likelihood ratio test.

#### Relationships between trial-wise computational estimates of learning and pain responses

To test our main hypothesis, we assessed the relationship between trial-wise PEs estimated using the winning RW model and four pain-related responses: pain intensity ratings, the amplitude of the N2-P2 component, gamma-band power and the amplitude of the NFR. We performed linear regressions between these measures at each trial within each participant and tested the significance of each relationship across participants using a two-tailed one-sample t-test on the slopes. We report results corrected for multiple comparisons using the Holm-Bonferroni procedure.

We additionally assessed if PEs modulated pain responses through the spinal modulation of nociceptive activity as measured by the NFR. To this end, we performed multilevel mediations with the NFR as a mediator of the relationship between PEs and the other pain responses. The mediation models were implemented with the CanLab mediation toolbox (<u>github.com/canlab/MediationToolbox</u>; [59,60]), and we assessed the significance of each path using a bootstrap resampling procedure with 10,000 bootstrap samples.

#### Relationship between pain responses and anticipatory EEG

We additionally assessed the relationship between anticipatory EEG responses to the conditioned cues and computational estimates of pain expectations (1 - PE), NFR amplitude and pain ratings. To this end, we used mass univariate analyses and cluster-based permutation tests [36] performed separately for the ERP and TFR data, with clusters being defined on the basis of temporal and spatial adjacency for ERP and temporal, spatial and spectral adjacency for TFR.

To perform these tests, the data at each channel was first z-scored across all time points and epochs for ERPs and all time points and epochs for ERPs frequencies between TFR within each participant. Ordinary least square linear regressions were then performed between each of the three z-scored variables and the z-scored ERP amplitude or the z-scored power within each participant at each time point and location or time point, location and for frequencies between 1-50 Hz. The beta values for each participant were then submitted to a two-tailed one-sample t-test comparing the group average beta value against 0 at each time point and location or time point, location and frequency, and the results were corrected using cluster-based permutations with 5000 permutations, a cluster entering the threshold of p < .01 two-tailed and a Bonferroni corrected cluster significance threshold for the three variables tested (0.05/3), two-tailed.

For the sake of completeness, we additionally performed similar analyses but without relying on the computational estimates of pain expectations. These model-free analyses compared the average ERP amplitude or TFR power across cue conditions. These analyses are described and reported in Supplementary Materials since they revealed similar results to the model-based analyses described above (see Supplementary Figure 2).

## Results

### Pain intensity ratings

On average, participants rated the electrical stimulations as 26.88 on the 0 to 100 visual analog scale (SD= 20.20, range: 0.70-68.17; Figure 1c). A repeated-measures ANOVA comparing average pain ratings across the six blocks indicated that there was no evidence of a change in pain perception across blocks [F(1.76, 52.95) = 0.29, p = 0.72].

### SCR and computational modelling

SCR amplitude at each trial served as an indicator of novelty and learned shock expectations (Figure 2a), displaying higher responses for new stimuli (CS-1) and stimuli previously associated with a shock (CS+). A linear mixed model verified that SCR varied among the four stimulus conditions (CS-1, CS-2, CS+, CS-E) [*χ*^2^ (3) = 233.38, p < 0.001]. This effect interacted with block number [*χ*^2^ (1) = 25.74, p < 0.001], indicating that the SCR response to CS+ was higher in block 1 than in other blocks [ *χ*^2^ (1) = 13.93, p < 0.001], while the other conditions did not vary across blocks [ps > 0.12].

**Figure 2.**
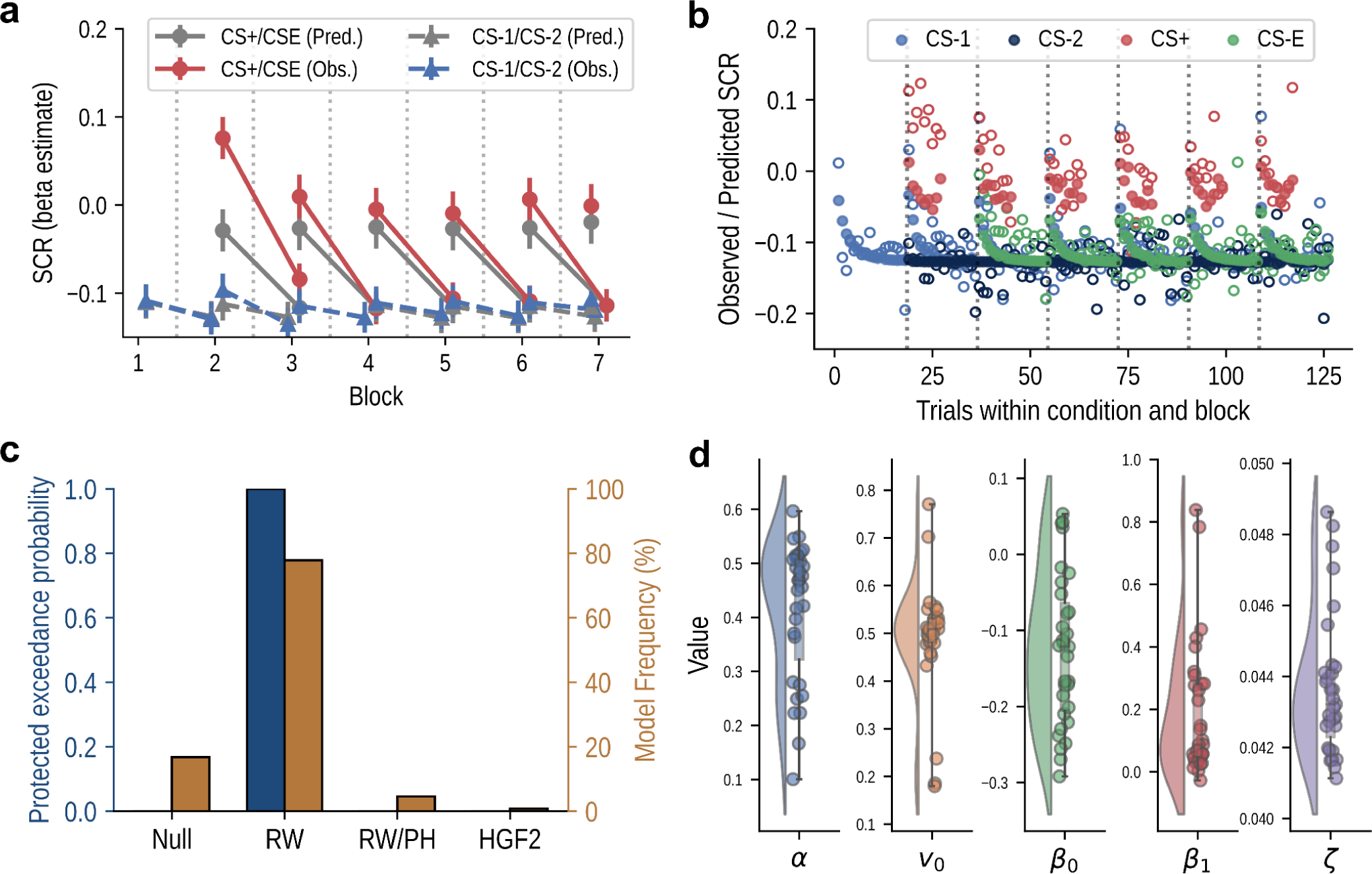
Skin conductance and computational model results. **(a)** Mean observed skin conductance response (Obs. colored) and mean predicted response (Pred. in gray) as a function of condition and experimental block. The lines between markers show the transition of one stimulus from one block to the next. Error bars show the standard error of the mean. **(b)** Observed (empty) and predicted (filled) SCR response for each trial. The dashed lines show the start of each experimental block. Note that reinforced trials were not predicted. **(c)** Results of the Bayesian model selection of the three inter-cue computational models showing the superiority of the Rescorla-Wagner model (RW) over a null model (no learning), as well as the combined RW/Pearce-Hall hybrid model (RW/PH) and the 2-level Hierarchical Gaussian Filter (HGF2) model. **(d)** Distributions of free parameters (see Methods) across participants for the winning RW model.

To quantify participants’ learning dynamics throughout the task, we evaluated the ability of various computational learning models to account for variations in single-trial SCR. The Bayesian model comparison revealed that the inter-cue family of models outperformed the simple models (Figure S1) and that the inter-cue RW model provided the best fit to the data in most participants (Figure 2c). The distributions of the free parameters across participants show that for most participants, the expectation was a predictor of the SCR response (Figure 2d). A visual inspection of the model’s predictions also supported a good fit to the data (Figures 2a and 2b).

### Prediction errors are associated with increased pain perception and physiological pain responses

This analysis of the relationship between trial-wise PEs estimated using the RW model and pain responses revealed a significant relationship between the prediction error and all of the pain responses (Figure 3b and Figure 3c), indicating that more surprising noxious stimuli led to increased physiological responses and were perceived as more intense. The multilevel mediations with the NFR as a mediator of the relationship between PEs and the other pain responses indicated that the NFR significantly mediated the relationship between PE and pain ratings and between PE and gamma power (Figure 3d). The mediation of the relationship between prediction error and N2-P2 amplitude by the NFR did not reach significance (Figure 3d).

**Figure 3.**
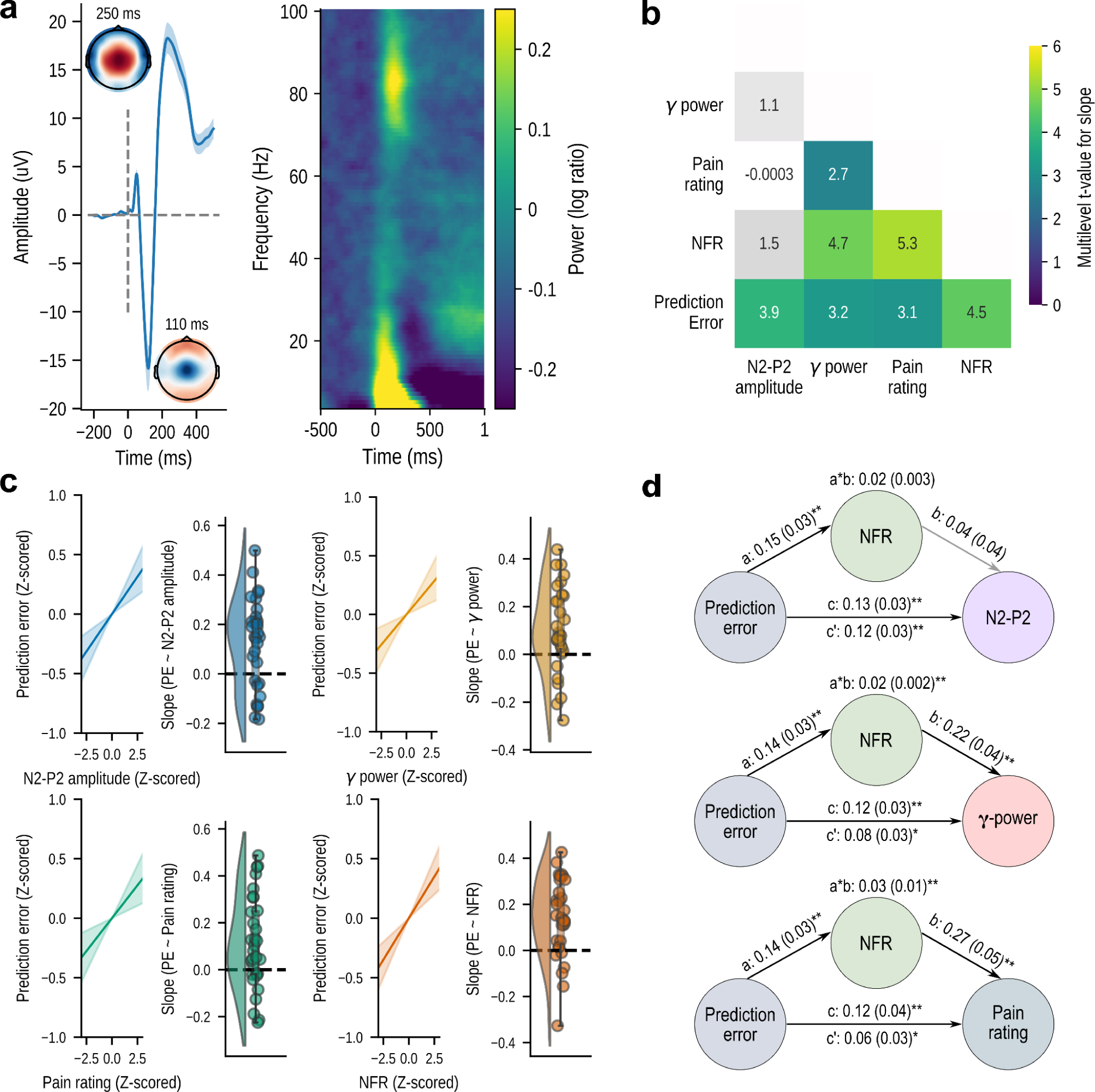
Pain responses and their relationship with prediction errors. **(a)** ERP response to the shock at Cz and topographic maps showing scalp distribution of the amplitude at each peak and TFR response to the shock at Cz showing the gamma band power. **(b)** Matrix showing the relationship between pain responses as the t-value for the second level test on the slopes of each participant. The colored squares show relationships surviving the Holm-Bonferroni corrected significance threshold and the gray squares show non-significant relationships. **(c)** Average and distribution of slopes across participants for the relationship of each pain response with prediction error. **(d)** Mediation of the relationships between prediction error and N2-P2 amplitude, gamma power and pain ratings by the NFR. *p < 0.05 **p < 0.005.

### Prediction errors are associated with increased anticipatory cortical responses

The amplitude of the late ERP response to pain predictive cues between 400 and 600 ms was closely related to the magnitude of the trial-wise computational estimate of pain expectation (Figure 4a). The mass univariate regression indicated that subjective pain expectation was significantly related to the ERP amplitude at parieto-occipital and central sites from 400 ms after the cue onset (Figure 4b). Pain expectations were also negatively related to anticipatory alpha-to-beta power. The results shown in Figures 4c and 4d reveal a significant negative relationship between pain expectation and the alpha and beta power at parieto-occipital sites and beta power at central sites.

**Figure 4.**
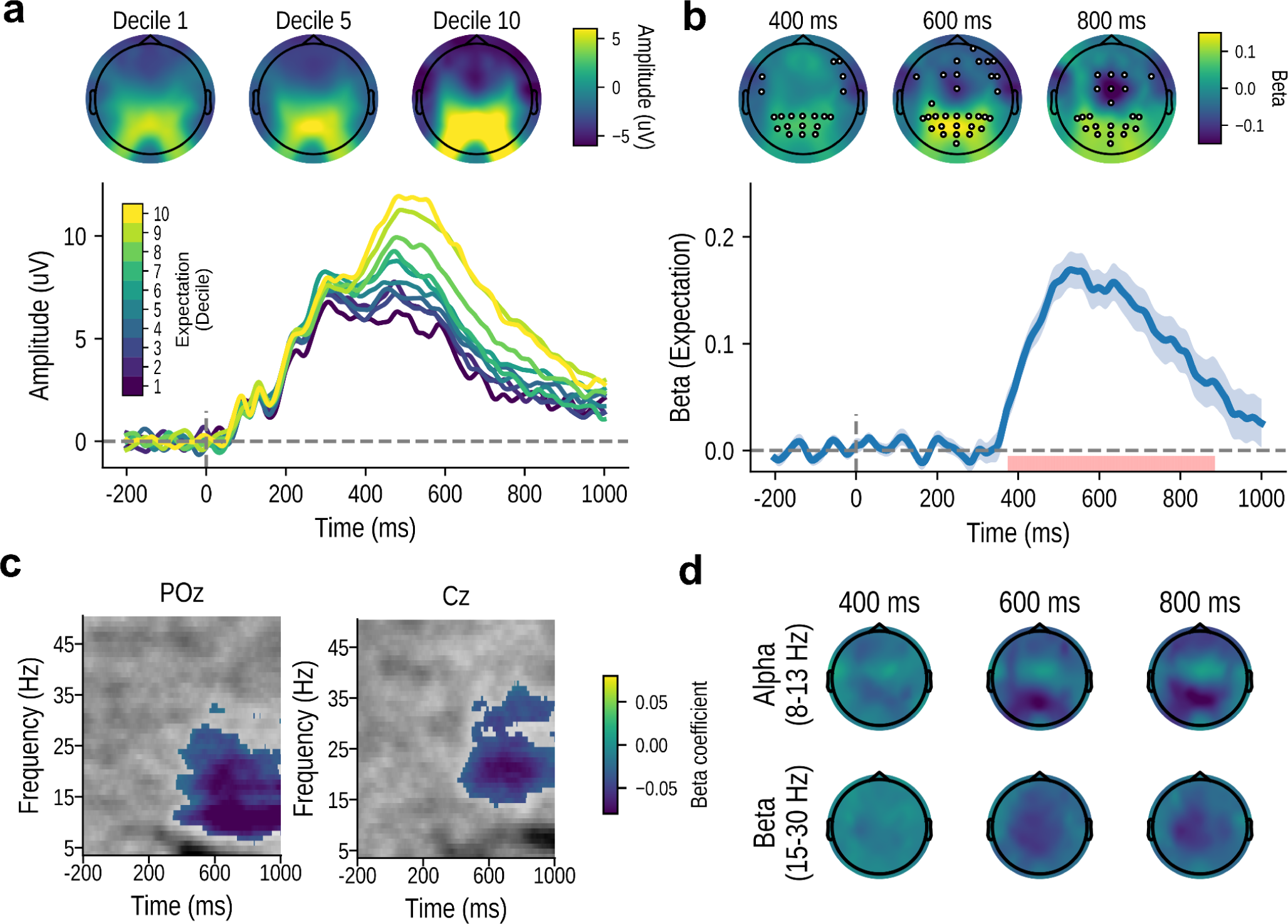
Results of the model-based analyses on the EEG response to the predictive cues. **(a)** Topographic map and ERP amplitude for various expectation levels at channel POz showing the increase in amplitude with the increase in pain expectation. **(b)** Beta estimates at POz (line plot) and at 400-800 ms (topographic plots) for the mass univariate regression of expectation on the ERP amplitude. The red lines and white circles show significant time points and locations (p < 0.05, cluster-based correction). **(c)** Beta coefficients for the mass univariate regression of pain expectation on TFR activity. The colored values show significant differences at p < 0.05, cluster-based correction. **(d)** Topographic maps illustrating the mean beta coefficients for the alpha and beta bands at different times.

While a positive relationship between the late ERP amplitude at parieto-occipital sites was also observed for the pain ratings and NFR, the statistical tests of these relationships did not reach significance, and no clear relationship was observed between these two variables and time-frequency power (Supplementary Figure 3).

## Discussion

One of pain’s primary functions is to teach us what may cause harm so that we can avoid pain in the future. This study sought to test the hypothesis that subjectively perceived pain is modulated at the spinal level by the informational value of nociceptive inputs. In support of that hypothesis, we found that subjective pain perception increases with the prediction error (PE) signals necessary to learn from surprising pain. Moreover, nociceptive flexion reflexes (NFRs) and EEG markers of cortical nociceptive processing (N2-P2 and gamma-band power) also increased with PEs, suggesting that prior expectations can influence the transmission of nociceptive signals from the spinal cord to the brain. Thus, nociceptive inputs to the brain appear to carry a PE signal that is used for learning to predict the probability of future pain, suggesting that perceived pain is profoundly tied to its learning and adaptive function.

Our experimental paradigm was designed to produce frequent PEs by repeatedly introducing new conditioned stimuli during the task. Expectations about pain and PEs were estimated on a trial-by-trial basis by fitting different candidate learning models to the conditioned SCRs [52,58]. This procedure indicated that a simple delta-rule Rescorla-Wagner model [46] was the best fit for our data compared to other more complex models taking into account uncertainty [39,44]. The reason why this simpler model provided a better fit could be because a large number of cues were used and regularly introduced during the task.

Consistent with our prior studies [56,57], models that incorporated information transfer from reinforced cues to others not presented yielded the best fit for our data. This inter-cue learning transfer may stem from the uniform reinforcement schedule in the tasks we used. It could also suggest a broader transfer of learning across different task blocks, an aspect not fully addressed by the models in our current study and which might influence the relationship between learning and pain perception. Future research should explore designs that enable explicit modelling of task structure. For instance, deliberate manipulation of reinforcement contingencies throughout the task to reduce the regularity of the reinforcement schedules [42] could provide valuable insights into how such manipulations influence pain perception.

Computational PE estimates derived from the computational model predicted all dependent variables recorded in response to electric shocks, including pain ratings, NFR amplitude, N2–P2 amplitude, and gamma-band power. These findings are particularly interesting given that the data used to estimate PEs (anticipatory SCR responses to unreinforced shocks) was independent from the responses to the shocks, supporting the notion that pain’s learning function coordinates psychophysiological responses in anticipation and response to pain. More specifically, modulation of NFR amplitude by PEs suggests that nociceptive signaling takes on a learning function as soon as nociceptive signals enter the central nervous system in the dorsal horn of the spinal cord. Moreover, correlations between NFR amplitude, PE estimates and EEG responses to pain also further support the interpretation that prediction errors do not only influence the excitability of spinal motoneurons, but also gate the transmission of nociceptive signals from the spinal cord to the cortex. This interpretation is further supported by a non-human electrophysiological study showing that neurons of the dorsal horn can reflect behaviorally relevant task signals and therefore do not only passively transmit sensory signals to the brain [16]. Overall, these findings provide converging evidence suggesting that the main function of subjectively experienced pain may be to indicate the need to adjust our behavior rather than simply representing the intensity of the nociceptive stimulus [9,31].

The relationships between PEs and the N2-P2 amplitude and gamma-band power suggest that expectations can also affect the cortical response to nociceptive stimulation. Reduced cortical responses to pain when pain is expected likely reflects a downstream effect of reduced spinal nociception [7,13, but see 27]. This interpretation was supported by the significant correlations between NFR amplitude and gamma-band power. However, correlations between NFR amplitude and N2-P2 amplitudes were not found to be significant. This could be explained by previous findings suggesting that the N2-P2 may not be specific to nociceptive input [40] but rather reflects general arousal and saliency. Still, N2-P2 amplitude correlated with PE estimates, suggesting that the non-significant positive association between N2-P2 amplitude and NFR amplitude might have reached significance with added statistical power. By contrast, previous research has suggested that gamma-band power may be a more specific measure of experienced pain [26,45] which may explain the strong correlations between gamma-band power, NFRs and subjective pain ratings. There is however a need for caution when interpreting the results of the relationship between gamma-band power and pain responses. Indeed, gamma-band power elicited by pain has been shown to share a similar topography to gamma-band activity related to facial movements and micro-saccades [10,49]. Although we employed ICA to attempt to eliminate muscle artifacts from the data, it is still possible that the results were partially influenced by residual muscle artifacts, potentially correlated with prediction errors.

Overall, these findings correspond well with the notion that conditioned responses serve to prepare the organism for the upcoming homeostatic disturbance and therefore reduce the need to react to the painful stimulus itself [53]. This way, pain perception is reduced when pain is expected and amplified when unexpected. However, these findings also conflict with prior studies showing that perceived pain and NFRs are reduced when the expected probability of pain is low [56,57]. In this set of studies, cue-outcome contingencies remained stable for a large number of trials such that anticipatory SCRs were better fitted by a hybrid Pearce/Hall learning model with an associability term. Associability increases following surprising events to facilitate learning when the environment is more uncertain. These studies found a positive relationship between associability and perceived pain and NFRs, and a negative relationship between PEs and pain and NFRs. By contrast, in the current experiment, associability was kept constantly high due to the frequent appearances of new cues, and we observed a positive relationship between pain and PEs. One potential explanation for the apparently discrepant findings is that pain may be positively related to PEs when associability is high because participants actively try to predict pain but are negatively related to PEs when associability is low because there is less of a need for learning. This idea is in line with previous demonstrations of the opposite effects of certain and uncertain expectations on pain perception and NFR amplitude [47].

Expectations of pain during the anticipatory period were shown to be associated with increased LPP amplitude at central and parieto-occipital locations and increased alpha/beta decrease in power at occipital and centro-parietal sites starting around 400 ms after cue onset. These findings largely replicated previous studies on anticipatory responses to predictable aversive stimuli [5,41,43,51]. The late positive potential is thought to reflect attentional allocation and motivational salience [13,22] and might have reflected the threat value of the conditioned stimuli. By contrast, alpha/beta decrease have been linked to sensory predictions during both the anticipation of pain [41,55] and other sensory events [1,6,54]. In the present experiment, alpha/beta power decreased with the expected probability of pain, indicating that it might have reflected top-down signaling of an upcoming sensory event.

The late positive potential and changes in oscillatory power anticipatory responses to the predictive cues were not related to pain ratings or to the NFR amplitude. This contrasts with previous work showing a positive relationship between anticipatory responses and pain perception [2]. However, although not significant, we also observed a positive relationship between pain ratings and prestimulus EEG response. Moreover, it should be noted that anticipatory EEG activity and psychophysiological responses to pain were all connected through computational estimates of pain expectations and prediction errors. This underlines the central role that learning has to play in pain processing and suggest that the direct relationship between anticipatory activity and responses to pain could have reached significance with added statistical power.

Results from this study should be replicated and extended in different contexts. One limitation of this study is the use of a repeated reinforcement schedule, which limits the variability of the nature of the cue-pain associations. Future studies should assess the relationship between the informational value of nociceptive inputs and pain perception in an environment in which the probability of receiving pain after specific cues changes over time, therefore creating more uncertainty [7,28]. Furthermore, in this study, participants passively experienced the pain stimulations and could not avoid them. It would be interesting to further assess the effect of the informational value of pain on pain perception when participants can perform choices based on the available information to avoid future pain.

In summary, we showed that computational estimates of learned pain expectations are associated with central anticipatory responses and pain perception. These results further strengthen the idea of a link between pain and learning and suggest that understanding the influence of learning mechanisms on pain could help us understand when and why pain perception is modulated in health and disease.

## Acknowledgements

MPC was supported by a fellowship from the Canadian Institutes of Health Research and a career award from the Fonds de Recherche Québec–Santé (FRQS) during this work. This work was supported by a Discovery Grants from the Natural Sciences and Engineering Research Council of Canada (NSERC) to PR (Grant RGPIN-2013-06799), PJ (Grant RGPIN-2017-06679) and MR (Grant RGPIN-2016-06682) and a Canada Research Chair (CRC) in Brain Imaging of Experimental and Chronic Pain to MR and CRC in Experimental Cognitive Neuroscience to PJ. DKN is supported by the Canada Research Chairs Program, the Canadian Institutes of Health Research and the NSERC. PAB was supported by a scholarship from the FRQS in partnership with Parkinson Canada.

The authors declare no conflict of interest.

## Code and data availability

Due to ethical constraints, we cannot openly share the raw data. BIDS formatted raw data are available on request. The code and instructions for reproducing all analyses and figures are available at https://github.com/mpcoll/eeg_painlearning_2024.

## Model free analyses of the EEG response to the pain predictive cues

We carried out pairwise comparisons between conditions to test if the average ERPs tracked 1) the threat acquisition (CS+ vs CS-1), the effects of extinction vs. habituation threat extinction over habituation to the CS [(CS-1 vs CS-2) vs (CS+ vs CS-E)) and 3) if threat memory remained during extinction (CS-E vs CS-2). Pairwise comparisons were performed by testing the difference between each pair of conditions against 0 at all time points and channels using two-tailed one-sample t-tests, corrected using cluster-based permutations with 5000 permutations, a cluster entering threshold of p < 0.01 and a Bonferroni corrected significance threshold of 0.05/3. Grand-average ERP waveforms indicated that a late-positive potential emerging around 400 ms at parieto-occipital sites clearly distinguished between the CS+ from the other stimuli (Figure S1a) and tracked the extinction of the CS+ (Figure S1b). Mass-univariate analyses (Figure S1c) confirmed that the amplitude at parieto-occipital sites showed significant effects of threat acquisition (CS+ vs CS-1), threat extinction [(CS-1 vs CS-2) vs (CS+ vs CS-E)] and threat memory (CS-2 vs CS-E) at parieto-occipital sites (Figure S1d).

Similarly to the ERPs analyses, we averaged the TFR across trials and took the difference between conditions to obtain the average TFR representing the threat. acquisition, extinction and memory. To reduce computational demands, we restricted the analyses between 4 and 50 Hz and −0.2 to 1 s around the onset of the cue. We compared the conditions using three two-tailed one-sample t-tests carried out at all frequency bins, channels and time points in these intervals. Results were corrected using a cluster-based permutation procedure with 5000 permutations, a cluster entering threshold of p < 0.01 and a Bonferroni corrected significance threshold of 0.05/3. The average power in each condition suggests that the CS+ cues lead to a greater suppression in the alpha and beta bands at parieto-occipital sites (Figures S1e and S1g). The statistical test of this difference (Figure S1f) confirmed that clusters showing significant suppression of the alpha and beta rhythms at parieto-occipital sites (Figure S1h) during threat acquisition (CS-1 vs CS+) and to a lesser extent during extinction [(CS-1 vs CS-2) vs (CS+ vs CS-E)]. For threat memory (CS-2 vs CS-E), significant differences were only present in the alpha band.

**Figure S1.**
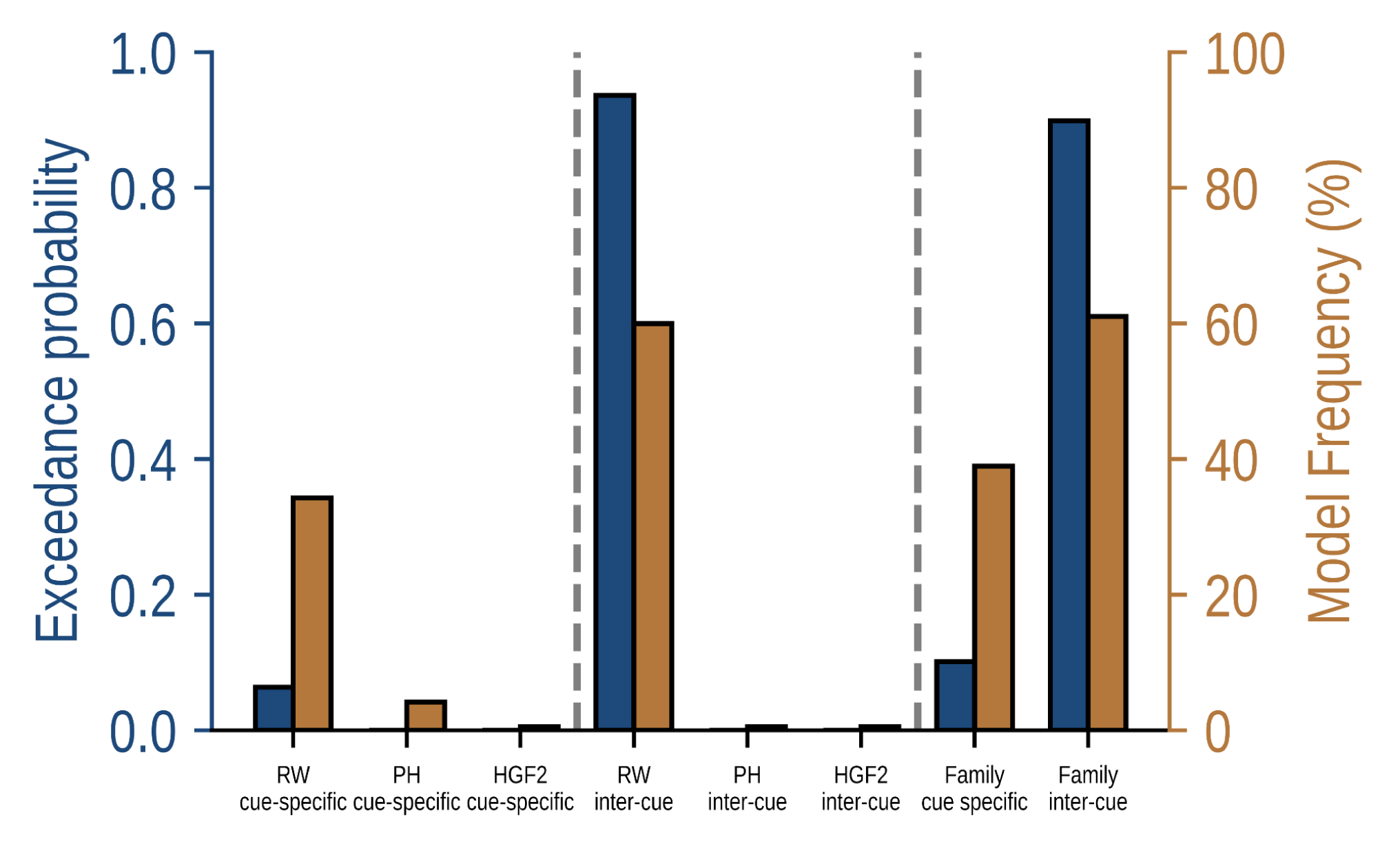
Bayesian model comparison for all computational models. Results of the Bayesian model selection comparing all models and the cue-specific and inter-cue families (see Methods).

**Figure S2.**
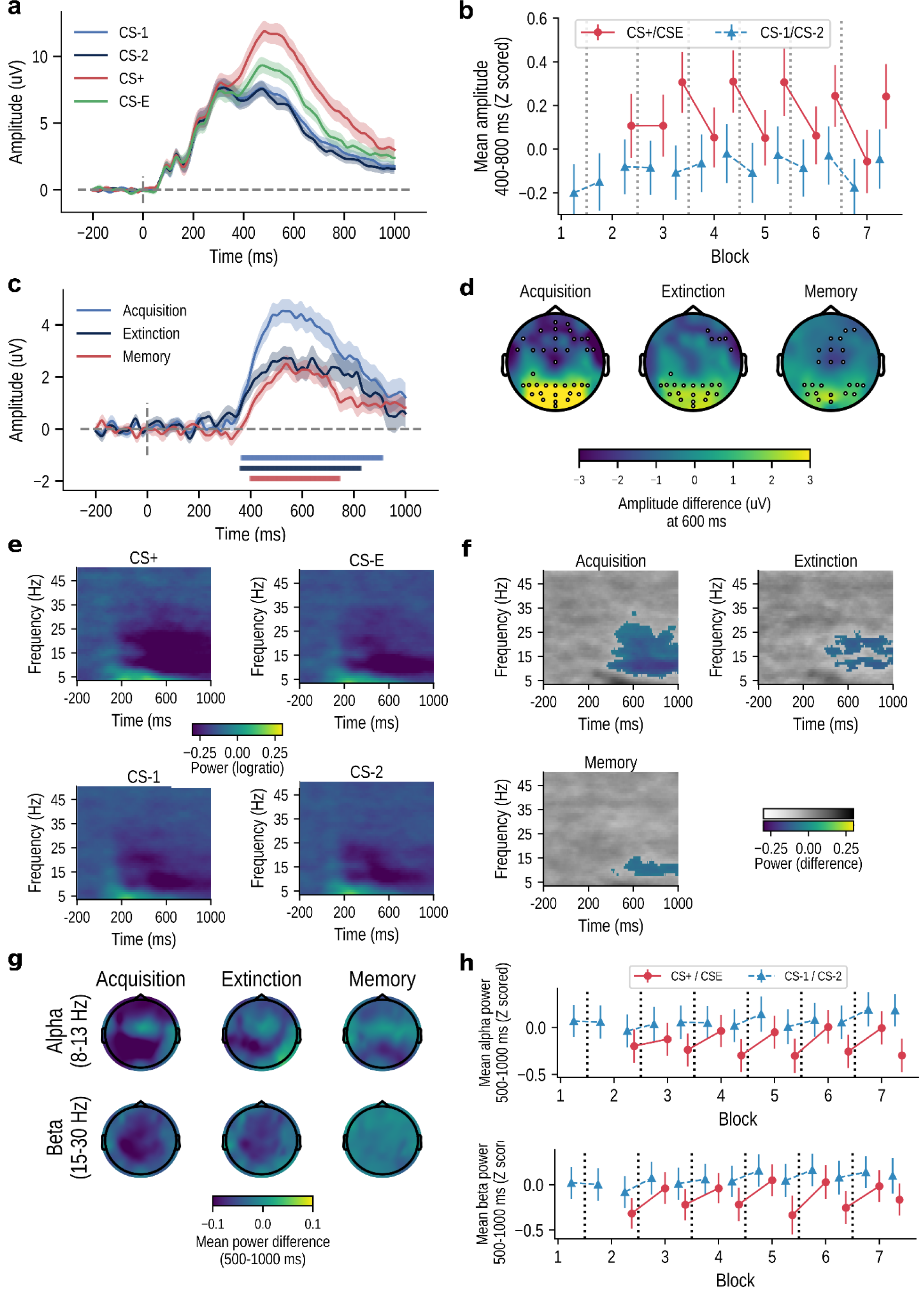
Results of the model-free analyses comparing EEG response to predictive cues as a function of condition. (a) Average event-related potential to the four cue conditions at POz. Shaded areas show the standard error of the mean. **(b)** Mean ERP amplitude between 400 and 800 ms at POz for each experimental block and cue. The lines between markers show the transition across blocks. Error bars show the standard error of the mean. **(c)** ERP differences between conditions at POz showing the threat acquisition (CS-1 vs CS+), extinction [(CS+ vs CS-E) vs (CS-1 vs CS-2)] and memory (CS-E vs CS-2). The shaded lines around the waves show the standard error of the mean and the shaded rectangles show time-points for which the difference was significantly different from 0 (0.05/3, cluster-based correction) **(d)** Scalp distribution of the difference between conditions. Highlighted channels show differences significantly different from 0 (0.05/3, cluster-based correction). **(e)** Mean power for each of the experimental conditions at POz. **(f)** Power differences between the conditions showing the threat acquisition (CS-1 vs CS+), extinction [(CS-1 vs CS-2) vs (CS+ vs CS-E)] and memory (CS-E vs CS-2) at channel POz. The colored values show significant differences at p < 0.05/3, cluster-based correction. **(g)** Topographic maps illustrating the mean power difference for each comparison in the alpha and beta bands between 500 and 1000 ms. **(h)** Mean power averaged between 500 and 1000 ms at POz for each experimental block and cue at the alpha (top) and beta (bottom) frequency bands. The lines between markers show the transition of one stimulus from one block to the next. Error bars show the standard error of the mean.

**Figure S3.**
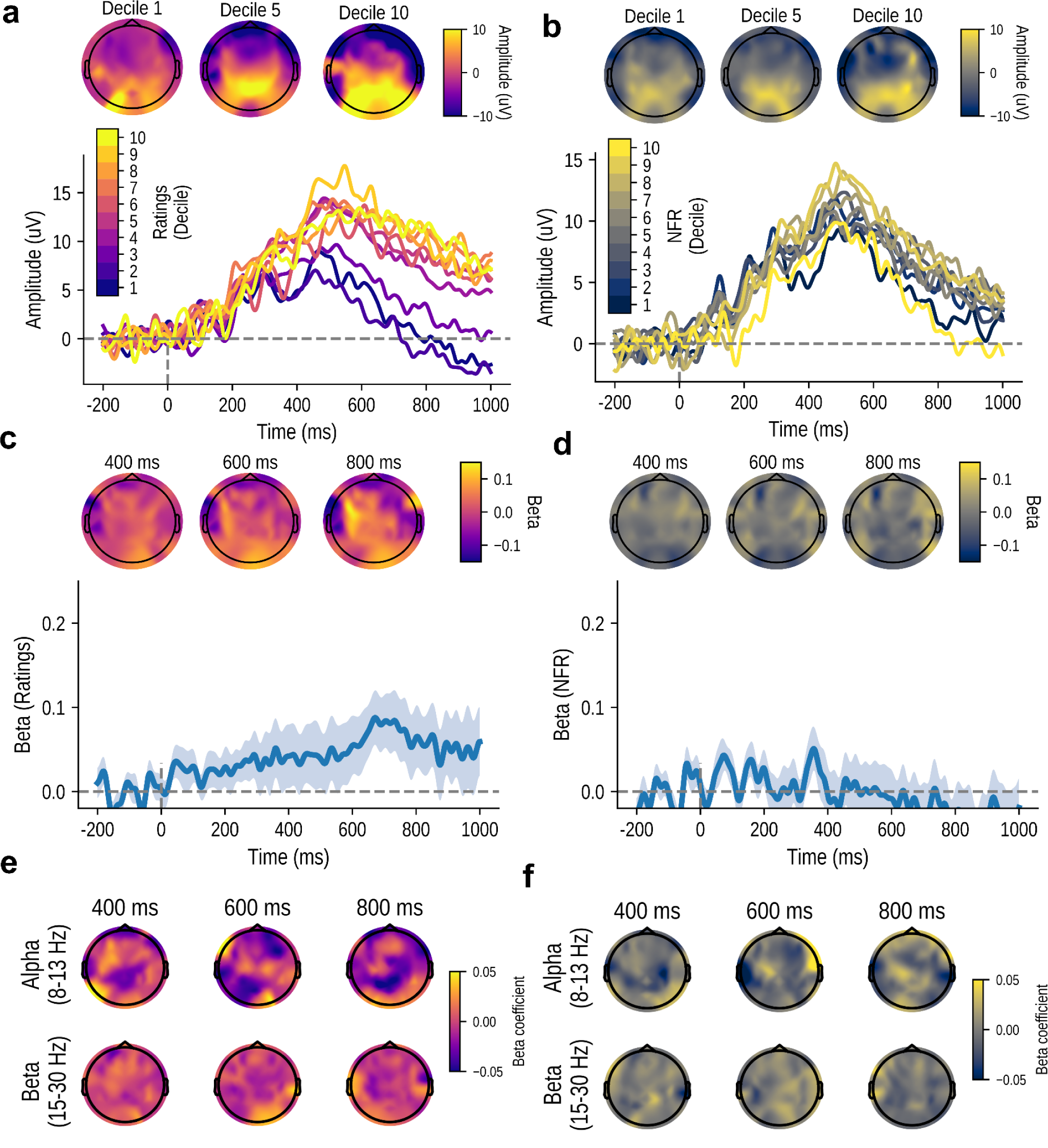
Relationship between pain ratings and NFR amplitude and EEG response to the predictive cues. **(a-b)** Topographic map and ERP amplitude for various levels at channel POz showing the differences in amplitude with the change in (a) subsequently perceived pain or (b) the NFR amplitude. **(c-d)** Beta estimates at POz (line plot) and at 400-800 ms (topographic plots) for the mass univariate regression of (c) pain ratings and (d) NFR amplitude on the ERP amplitude. **(e-f)** Topographic maps illustrating the mean beta coefficients for the alpha and beta bands at different times for the mass univariate regression of (e) pain ratings and (f) NFR amplitude on time-frequency power in response to the cues.

## Notes

### Competing Interest Statement

The authors have declared no competing interest.

### Summary of Updates

All results and figures updated following new analyses.

