## Supplementary Materials for "Pain reflects the informational value of nociceptive inputs"

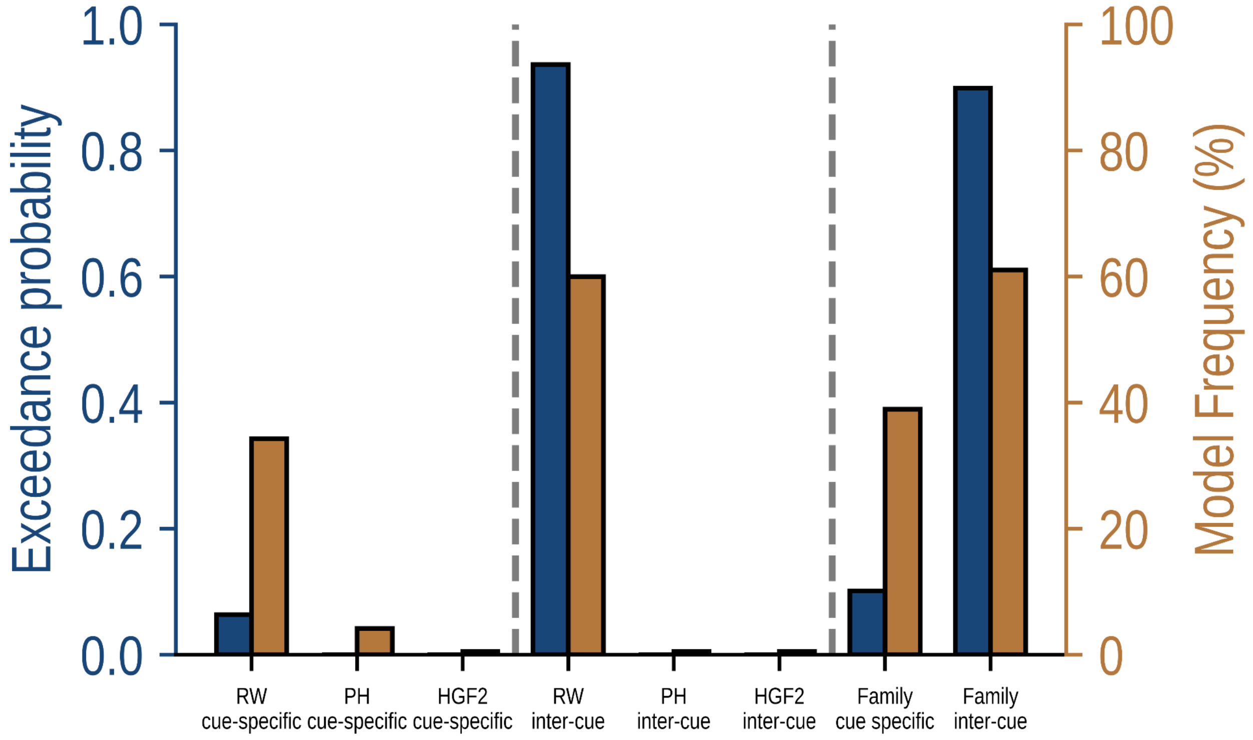


**Figure S1. Bayesian model comparison for all computational models.** Results of the Bayesian model selection comparing all models and the cue-specific and inter-cue families (see Methods).

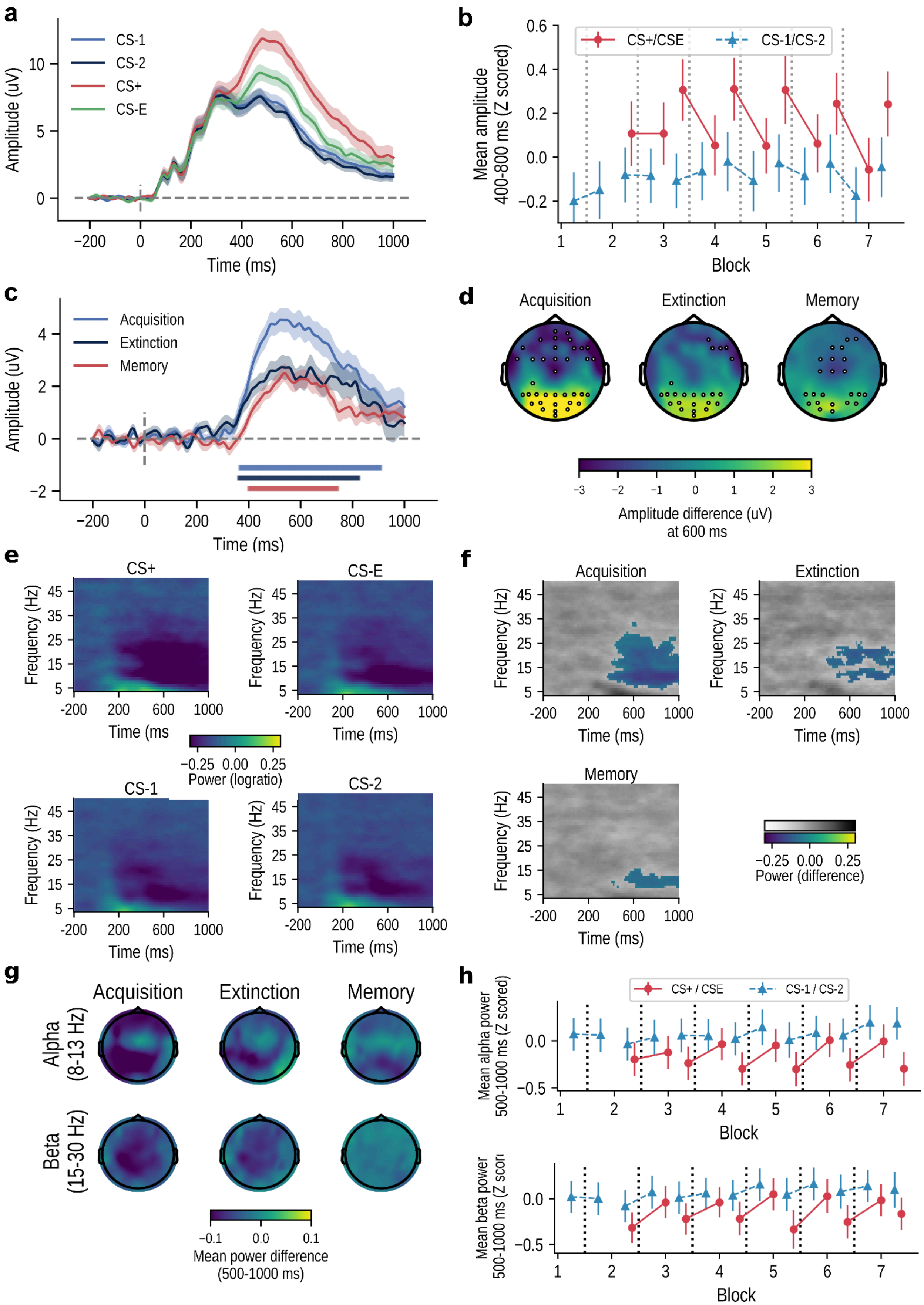
a
CS-1
CS-2
CS+
CS-E
Amplitude (uV)
Time (ms)
b
CS+/CSE
CS-1/CS-2
Mean amplitude
400-800 ms (Z scored)
Block
c
Acquisition
Extinction
Memory
Amplitude (uV)
Time (ms)
d
Acquisition
Extinction
Memory
Amplitude difference (uV)
at 600 ms
e
CS+
CS-E
CS-1
CS-2
Frequency (Hz)
Time (ms
Time (ms)
Power (logratio)
f
Acquisition
Extinction
Memory
Frequency (Hz)
Time (ms)
Power (difference)
g
Acquisition
Extinction
Memory
Alpha
(8-13 Hz)
Beta
(15-30 Hz)
Mean power difference
(500-1000 ms)
h
CS+ / CSE
CS-1 / CS-2
Mean alpha power
500-1000 ms (Z scored)
Block
Mean beta power
500-1000 ms (Z scor
Block

**Figure S2. Results of the model-free analyses comparing EEG response to predictive cues as a function of condition. (a)** Average event-related potential to the four cue conditions at POz. Shaded areas show the standard error of the mean. **(b)** Mean ERP amplitude between 400 and 800 ms at POz for each experimental block and cue. The lines between markers show the transition across blocks. Error bars show the standard error of the mean. **(c)** ERP differences between conditions at POz showing the threat acquisition (CS-1 vs CS+), extinction [(CS+ vs CS-E) vs (CS-1 vs CS-2)] and memory (CS-E vs CS-2). The shaded lines around the waves show the standard error of the mean and the shaded rectangles show time-points for which the difference was significantly different from 0 (0.05/3, cluster-based correction) **(d)** Scalp distribution of the difference between conditions. Highlighted channels show differences significantly different from 0 (0.05/3, cluster-based correction). **(e)** Mean power for each of the experimental conditions at POz. **(f)** Power differences between the conditions showing the threat acquisition (CS-1 vs CS+), extinction [(CS-1 vs CS-2) vs (CS+ vs CS-E)] and memory (CS-E vs CS-2) at channel POz. The colored values show significant differences at $p < 0.05/3$, cluster-based correction. **(g)** Topographic maps illustrating the mean power difference for each comparison in the alpha and beta bands between 500 and 1000 ms. **(h)** Mean power averaged between 500 and 1000 ms at POz for each experimental block and cue at the alpha (top) and beta (bottom) frequency bands. The lines between markers show the transition of one stimulus from one block to the next. Error bars show the standard error of the mean.

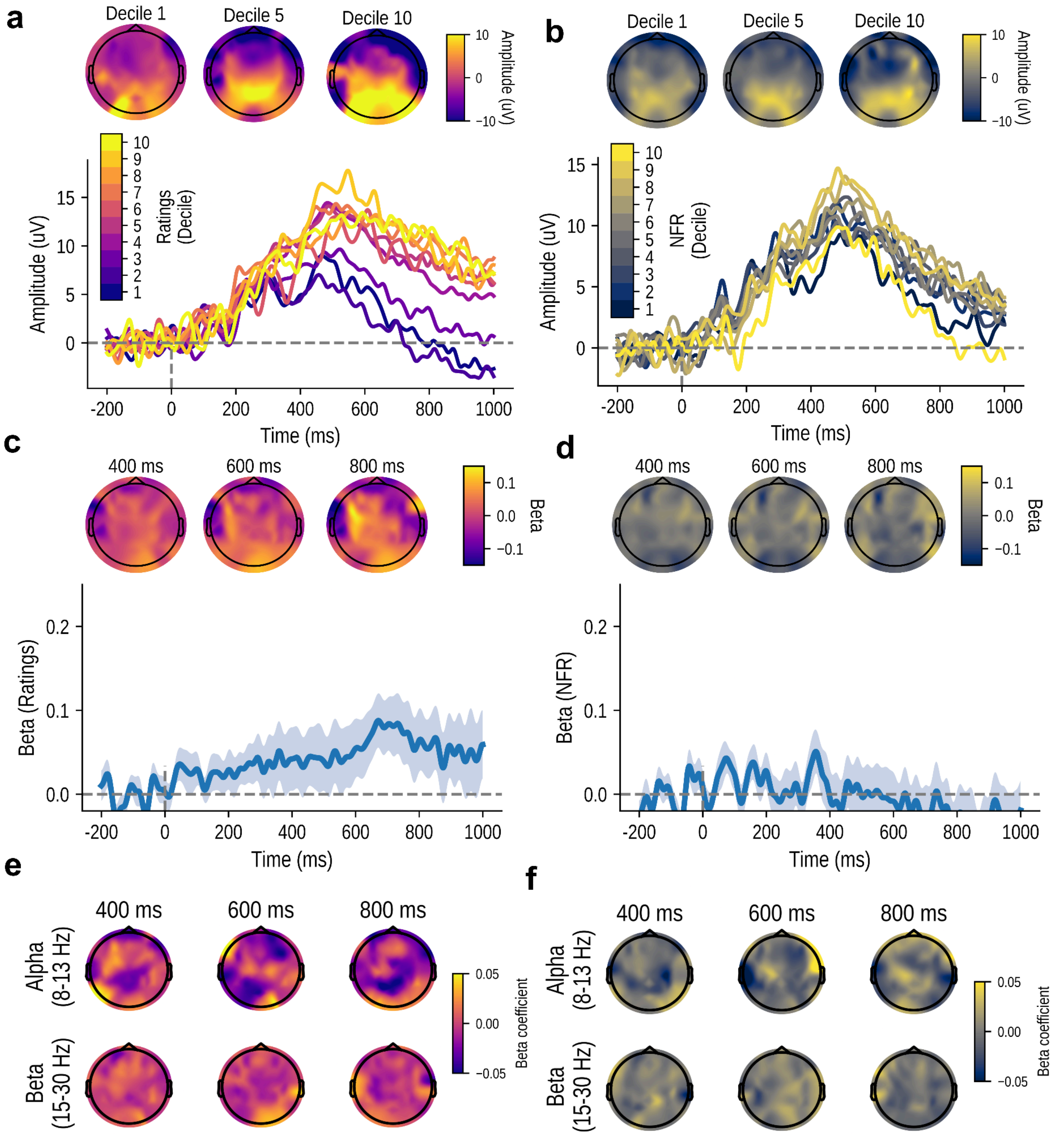


**Figure S3. Relationship between pain ratings and NFR amplitude and EEG response to the predictive cues. (a-b)** Topographic map and ERP amplitude for various levels at channel POz showing the differences in amplitude with the change in (a) subsequently perceived pain or (b) the NFR amplitude. **(c-d)** Beta estimates at POz (line plot) and at 400-800 ms (topographic plots) for the mass univariate regression of (c) pain ratings and (d) NFR amplitude on the ERP amplitude. **(e-f)** Topographic maps illustrating the mean beta coefficients for the alpha and beta bands at different times for the mass univariate regression of (e) pain ratings and (f) NFR amplitude on time-frequency power in response to the cues.
